# Motor mirror neurons may not be a predictor of learning but reflect the effect of motor learning

**DOI:** 10.1101/2023.12.16.572003

**Authors:** Pomelova Ekaterina, Feurra Matteo, Popyvanova Alena, Nikulin Vadim, Solodkov Roman, Banjevich Tamara, Blagovechtchenski Evgeny

## Abstract

The mirror neurons system (MNS) fires during both the performance of an action and observation of the same action being performed by another. On the level of motor output, activation of the MNS is thought to be represented in the phenomenon of motor resonance, which manifests in a muscle-specific increase in corticospinal excitability during action observation. This study focused on how and to what extent sensorimotor learning alters the initial mirror response and whether the rate of sensorimotor learning is associated with pretraining or post-training levels of mirror response. The study involved 23 healthy adults aged 22.7 years. The experiment consisted of six sessions. On the first and last days, a transcranial magnetic stimulation session was held to assess the putative activity of mirror neurons, as reflected in the level of motor-evoked potential facilitation during action observation in different conditions. From the second to the fifth sessions (four sessions in total), the sensorimotor learning part was performed, as represented in the form of a serial reaction time (SRT) task. We observed a statistically significant decrease of reaction time in the process of learning in the SRT task and motor facilitation during action observation, thus reflecting the process of putative mirror neurons’ activity. However, our data demonstrate that the sensorimotor learning rate was not associated with either pretraining or post-training estimates of motor facilitation during action observation and that sensorimotor learning does not affect the pattern of motor resonance.

## Introduction

Although the initial discovery of mirror neurons was made with the help of single-unit recording in the macaque’s premotor cortex (area F5) (1), the existence of an analogous system in humans is currently supported by the solid body of evidence (2,3). Despite the previous assumption that the mirror response is an innate capability of humans to represent others’ actions in one’s own motor code (4), more recent theories emphasize the crucial role of sensorimotor learning in the development of this phenomenon (5–7). On the level of motor output, activation of the mirror neurons system (MNS) is thought to be represented in the phenomenon of motor resonance, which manifests in a muscle-specific increase of corticospinal excitability during the action observation of corresponding effectors (8). Because mirror neurons are a reflection of the activity of the external environment and our adaptation to the external environment is manifested in learning, one can assume that the activity of mirror neurons may serve as a predictor of the success of the learned movement.

Associative sequence learning (ASL) theory states that the initially separated motor and visual representations of an action become coupled in the process of congruent sensorimotor learning, during which activations in motor neuronal populations coincide with activations in visual neuronal populations, thus promoting the emergence of visuomotor associations based on the principle of Hebbian learning (9–11). Ideomotor theory considers sensorimotor learning as the key process in the development of mirror response as well, complementing ASL theory by suggesting that actions are additionally represented in the form of the predicted sensory consequences of these actions (12). Although these theories have certain conceptual differences, mainly restricted by the assumption of ideomotor representations’ existence or absence thereof, they are consistent with each other when it comes to the emphasized role of sensorimotor learning in the development of motor mirroring mechanisms (13). However, despite the data available in the literature, the question of whether the process of mirroring is necessary for learning or instead represents the process of learning reflecting, for instance, interactions with the external world and attentional processes remains open.

The present study focused on how and to what extent sensorimotor learning alters the initial mirror response. Moreover, it is important to assess whether the rate of sensorimotor learning is associated with pretraining or post-training levels of mirror response. The following hypotheses were formulated: First, we expected to find a correlation between the rate of sensorimotor learning and the facilitation of motor excitability during action observation in the pretraining session. Second, we expected to observe a change in the pattern of motor resonance in the post-training session compared with the pretraining session, which would demonstrate a causal effect of the sensorimotor learning process on the motor resonance phenomenon.

## Materials and methods

### Participants

Overall, 24 participants participated. The participants were recruited two years ago (01/2020 -12/2020) from social media. One participant missed one day of the behavioral part of the experiment, and thus, their data were not included in the analysis. Therefore, we have data from 23 participants (16 females, 7 males) available for analysis. The mean age of the participants was 22.7 years (SD = 2.18). All participants were right-handed, with normal or corrected to normal eye sight, and they had no history of neurological diseases or traumas. Each participant signed the informant consent form before the start of the experiment.

### Time course of the experiment

For each participant, the experiment consisted of six sessions. All sessions were held consecutively on separate days. On the first day, a pretraining transcranial magnetic stimulation (TMS) session was held to assess the putative activity of mirror neurons, as reflected in the level of motor evoked potentials (MEPs) facilitation during action observation in different conditions (Baseline, Baseline Hand, Pinch, Button, and Baseline Post; see the description below). These five conditions were implemented during the stimulation of the dominant or nondominant hemispheres (and simultaneous recording of the electromyography (EMG) in the contralateral hand), resulting in 10 conditions being applied during the pretraining TMS session. From the second to the fifth session (4 sessions in total), the sensorimotor learning part was performed, as represented in the form of a serial reaction time (SRT) task. The task was implemented separately for both hands (dominant and nondominant). On the sixth day, a post-training TMS session, which was completely identical to the pretraining TMS session, was held to assess changes in the pattern of motor resonance.

### TMS session

TMS was applied over the left or right primary motor cortices (M1) through a MagPro X100 (MagVenture) stimulator with a C-B60 Butterfly induction coil (MagVenture) with biphasic pulses. For neuronavigation purposes, the MNI template was adjusted to the participants’ individual coordinates with the help of a TMS navigation system (Localite TMS Navigator, Localite GmbH). This approach ensured consistency of the defined hotspot throughout the experiment and across the sessions. The coil was held tangential to the scalp, with the handle pointing backward and laterally, at a 45° angle from the midline sagittal axis of the participant’s head. Single TMS pulses were delivered to find the optimal hotspot for the first dorsal interosseous (FDI) muscle (i.e., the scalp point, stimulation of which at threshold intensity elicited MEPs in the FDI muscle of the contralateral hand) (14). The resting motor threshold (RMT) for the given hotspot was determined as the minimal stimulation intensity inducing MEPs of minimum 50 µV in a resting muscle in 5 out of 10 given pulses (15). For experimental purposes, a stimulus intensity of 120% from RMT (120% RMT) was used. For each condition, 20–30 MEPs were recorded. Before each pulse, the “Go” command was voiced to prepare participants for the focused observation of the action executed. In the case of action observation, pulses were delivered in the middle or final phase of the action. This variability in the timing of the stimulation was introduced to prevent participants from predicting the exact moment of the stimulus application.

The following five conditions were implemented:

1. Preobservation resting state (baseline), during which the participants were instructed to sit relaxed with eyes open
2. Observation of static hand (Baseline Hand), during which the participants were asked to observe the static hand of the actor (Fig 1a.1)
3. Observation of pinch-to-grip movement (Pinch), during which the participants observed the actor who executed the pinch-to-grip movement (Fig 1a.2)
4. Observation of button-pressing movement (Button), during which the participants were asked to observe the actors’ hand pressing the button on the keypad (Fig 1a.3)
5. Postobservation resting state (Baseline Post), which was the same as the preobservation resting state but was recorded at the end of the experimental session.

**Fig 1.**
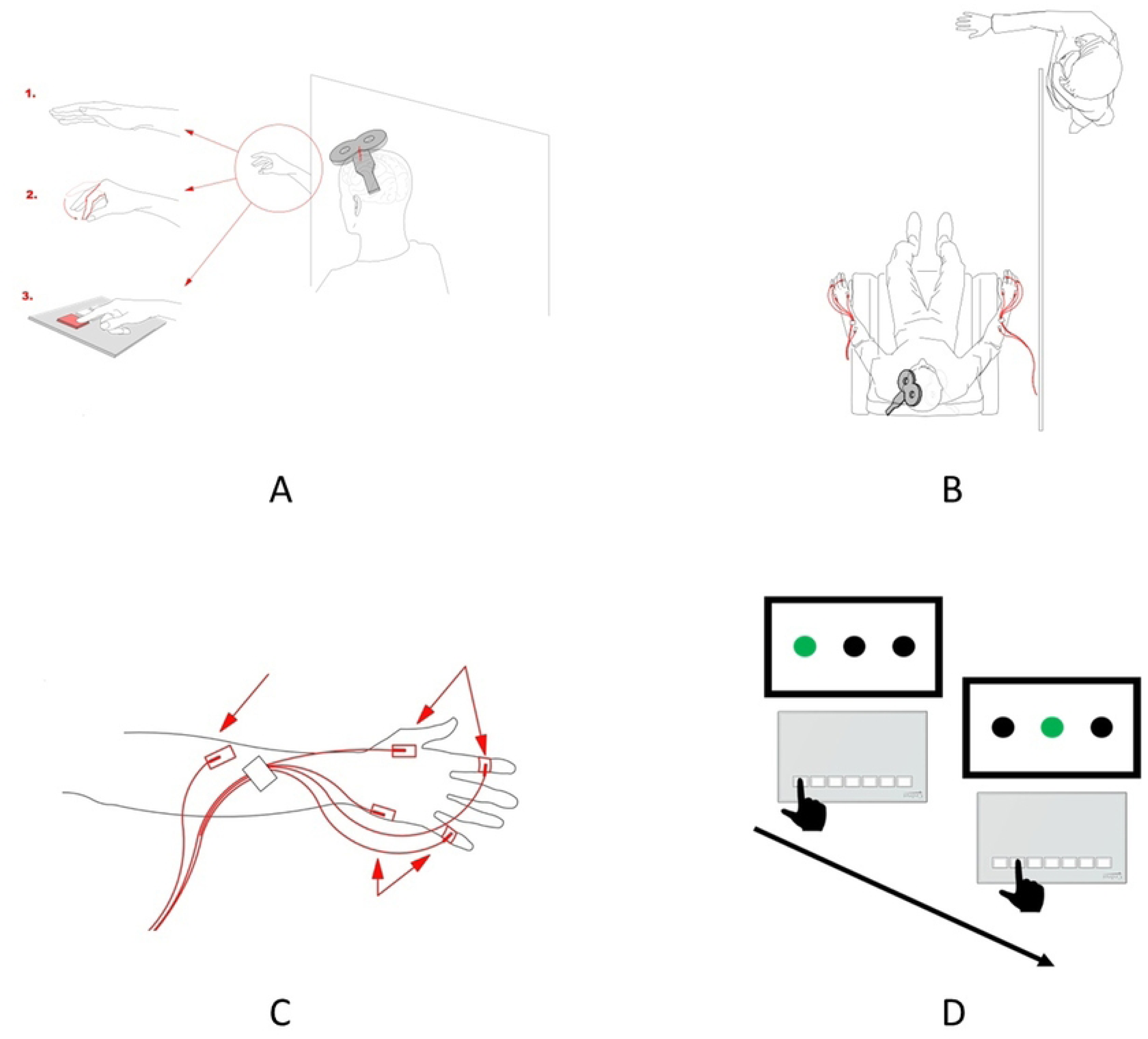
A) Schematic representation of the experimental conditions implemented during the TMS session. Observation of the static hand (1), observation of the pinch-to-grip movement (2), and observation of the button-pressing movement (3). In the preobservation and postobservation baseline conditions, the participants were requested to relax and observe the hand in front of them. B) Schematic representation of the spatial setup of the experiment. The participant can observe only the hand of the actor, while the rest of the actor’s body is located behind the curtain and, thus, is invisible to the participant. Red lines depict the location and placement of the EMG electrodes.C) Scheme of the surface electrode placement. Two electrodes were used for recording EMGs in the FDI, two electrodes were used for recording EMGs in the ADM, and the ground electrode was placed on the forearm. D) Two sequential trials of the SRT task. The black hand depicts the correct button-pressing movement in the trial—one of the three buttons, which corresponds to the green circle on the screen, should be pressed.

The order of Baseline, Baseline Hand, and Baseline Postconditions was fixed (because it is presented in the list above) to avoid possible long-term excitatory effects of observational conditions on the resting state and control conditions. Two observational conditions, on the other hand, were counterbalanced. There was a rationale behind including these particular conditions. A preobservation baseline was included to assess corticospinal excitability in the resting state. Postobservation baseline recording was added to ensure the absence of major changes in participant’s corticospinal excitability after the observation-related conditions. Additionally, the observation of the static hand was included to control for possible resting state differences because of the mere observation of a steady hand. The observation of the pinch-to-grip and button-pressing movements were expected to induce mirror-related muscle-specific facilitation of corticospinal excitability, thus allowing us to measure motor resonance as an index of mirror neurons’ activity. Both of these actions involve FDI, but not abductor digiti minimi (ADM); thus, we expected to observe an increase in peak-to-peak MEP amplitude, mainly for FDI. The difference between these two conditions is that pinch-to-grip movement was observed only during the TMS sessions and was not practiced during the behavioral part of the experiment, while the button-pressing movement was practiced extensively by participants during the completion of the SRT task.

All five conditions were implemented for both hemispheres and corresponding hands (dominant and nondominant). The participants observed only the actor’s hand (the rest of the body was located behind a curtain), executing actions with the hand corresponding to the one recorded by the participant (Fig 1 b). For both sessions (pretraining and post-training), experimental procedures were the same. To rule out the possible confounding factor of the changed hotspot, the same stimulation spot was used in the pretraining and post-training sessions, and consistency was ensured by saving navigational markers from the first session and using them during the second session.

### Electromyography recording

Before the application of the surface electrodes, participants’ skin on the desired areas was cleaned with ethanol (96% solution) and special scrub gel (Nuprep Skin Prep Gel, Weaver, and Company). For EMG recording, surface electrodes (Resting EKG Electrode, 3M Red Dot) were placed on the FDI and ADM muscles, and reference electrodes were placed on the joints of the index and little finger, respectively. The ground electrode was placed proximally on the hand. Thus, the overall electrode ensemble consisted of five surface EMG electrodes (Fig 1c). FDI was used as a target muscle because the implemented actions (pinch-to-grip and button-pressing movements) involved this muscle. The ADM muscle was used as a control muscle because the observed actions did not involve this muscle. The EMG was recorded with an ExG AUX box and BrainVision Recorder software installed on the computer.

For the EMG recording, the sampling frequency was set at 5000 Hz, and the resolution of the signal was set at 0.1 μV. EMG signals were filtered at a hi-pass filter of 10 Hz and a notch filter of 50 Hz (BrainAmp ExG amplifier). Triggers were sent from the TMS stimulator to the BrainVision software to extract epochs of the EMG recordings where TMS pulses were applied. Preprocessing of the EMG data was carried out with the help of MATLAB software and the “Berlin Brain–Computer Interface” toolbox.

### Serial reaction time task

The participants underwent sensorimotor training for four days in a row. On each day, the SRT task was performed for 10 minutes for each hand (dominant and nondominant), that is, 20 minutes per day in total. The order of task completion regarding the dominant and nondominant hands was counterbalanced across participants. Each participant had two days when the first hand was the dominant one and two days when the first hand was the nondominant one. For the SRT task, the Cedrus RB-740 response pad was used. The task itself was the following: participants were seated in front of the computer screen. On the screen, three black circles located horizontally were presented. When one of the circles lit up with green, the participant was instructed to press one of three buttons on the keypad, which corresponded to the green circle (Fig 1d). If the participant pressed the wrong button, the next trial was presented. The hand of the participant and keypad were positioned in such a way that pressing the button would require the involvement of the FDI muscle, but not the movement of the whole palm. The reaction time (RT) was recorded as the main dependent variable to assess learning pattern. The code for the SRT task was implemented with the Presentation software.

## Data analysis

### Learning estimation

First, behavioral data from the SRT task was analyzed because, based on these data, we could infer whether our task was sufficient for inducing sensorimotor learning in the first place. The main dependent variable in this part was the RTs of pressing the buttons on the keypad. The independent variables were Day (1, 2, 3, or 4) and Hand (Dominant or Nondominant). To investigate the dynamics of sensorimotor learning and check whether it did occur, we assessed changes in RTs across four days of training, while for the assessment of possible intermanual asymmetry, we had to measure the differences in RTs between two hands. Moreover, to investigate possible differences in the dynamics of sensorimotor learning between the two hands, we had to assess the interplay between our two independent variables. Accordingly, a two-way repeated measures ANOVA was performed on the acquired data: 4 (Days: 1, 2, 3, or 4) x 2 (Hand: Dominant or Nondominant).

### MEP extraction and estimation

As for the EMG data, peak-to-peak amplitudes of MEPs were used as an index of corticospinal excitability and, thus, as our main dependent variable for this part of the study. In the preprocessing stage, which was carried out with the help of the “Berlin Brain–Computer Interface” toolbox in MATLAB, high-pass (15 Hz) and notch (50 Hz) Butterworth filters were applied. Afterwards, epochs containing TMS-induced MEPs were extracted. Finally, the peak-to-peak amplitudes of MEPs were measured for both recorded muscles. In the next stage, statistical analysis of the acquired data was performed. Our independent variables at this stage were the following: Condition (Baseline, Baseline Hand, Pinch, Button or Baseline Post), Session (Pretraining or Post-training), and Hemisphere (Dominant or Nondominant). MEPs in the baseline and baseline postconditions were compared to rule out possible changes in general corticospinal excitability during the TMS session. For this, three-way repeated measures ANOVA was implemented: 2 (Condition: Baseline or Baseline Post) x 2 (Session: Pretraining or Post-training) x 2 (Hemisphere: Dominant or Nondominant). Facilitation of corticospinal excitability, which was used as an index of motor resonance, was calculated as the ratio of MEP amplitude in each condition to MEP amplitude in the “Baseline” condition (collapsed or not collapsed with the “Baseline Post” condition, depending on the result of baseline comparison; see below) in percent. Starting from this stage, after normalization, the ratio of MEP amplitude of each condition was used as the dependent variable and was used as the main dependent variable. To assess the differences in the change of corticospinal excitability, a three-way repeated measures ANOVA was performed on the data acquired from the FDI muscle: 3 (Condition: Baseline Hand, Pinch, or Button) x 2 (Session: Pretraining or Post-training) x 2 (Hemisphere: Dominant or Nondominant). To check the muscle specificity of the observed effects, the same analysis was performed on the data from the ADM muscle. For consequent post hoc analysis, a paired-samples t-test with Holm– Bonferroni correction was implemented.

### Statistics

Assessment of associations between the sensorimotor learning rate and MEP facilitation on the pre-training and post-training sessions was implemented with the Spearman’s correlation. The learning rate was normalized as the ratio of mean RT on the first day to mean RT on the fourth day for each hand. This particular ratio was chosen for the sake of comprehensibility: the higher is this ratio – the more pronounced the learning is. Comparisons were made between the levels of motor facilitation (for both sessions in both hands during observation of pinch-to-grip and button-pressing movement – 8 conditions) and learning rates (in dominant and non-dominant hands – 2 conditions), resulting in 16 comparisons in total. Multiple comparisons issue was accounted for with the help of Bonferroni correction. All statistical analysis, data handling and data visualization were carried out in the RStudio software. For the statistical analysis itself, “ez”, “corrplot” and “multcomp” packages were implemented. Data handling was mostly performed with “data.table” package. Finally, for data visualization, the “ggplot2” package was used.

## Results

### Learning estimation

Regarding the analysis of the SRT task, two-way repeated measures ANOVA ((Days: 1, 2, 3, or 4) x (Hands: Dominant or Nondominant)) was performed on the RT data. Mauchly’s test indicated that the assumption of sphericity was violated for the Day variable (W = 0.48, p = .009) and the interaction of the Day and Hand variables (W = 0.19, p < .0001). Thus, degrees of freedom were corrected using the Huynh–Feldt estimates of sphericity for Day (e = .74) and the interaction of Day and Hand (e = .55). A two-way ANOVA revealed significant main effects of Day (F (2.22, 48.80) = 91.59, p < .0001, h^2^ = .309) and Hand (F (1.22) = 5.62, p = .027, h^2^ = .011), while the interaction effect (F (1.65, 36.21) = 0.74, p = .530, h^2^ = .002) was not significant. Based on this pattern of effects, the performance was different on each day, with a gradual decrease in RTs across the sessions (Fig 2a), and this, in turn, suggests that our experimental manipulation was successful, hence inducing a pronounced sensorimotor learning effect in participants. Moreover, the significant main effect of the hand indicates that performance in the SRT task was generally better for the dominant hand compared with the nondominant one. However, the absence of an interaction effect signifies that the learning rate, which has been expressed in a gradual decrease in RTs across experimental sessions, did not differ between the dominant and nondominant hands. Additionally, for 11 subjects, accuracy was measured by calculating the ratio of the correct responses to all responses. Mean accuracy was equal to 95.4% (SD = 1.1 %), indicating a high level of hit rate for the implemented task.

**Fig 2.**
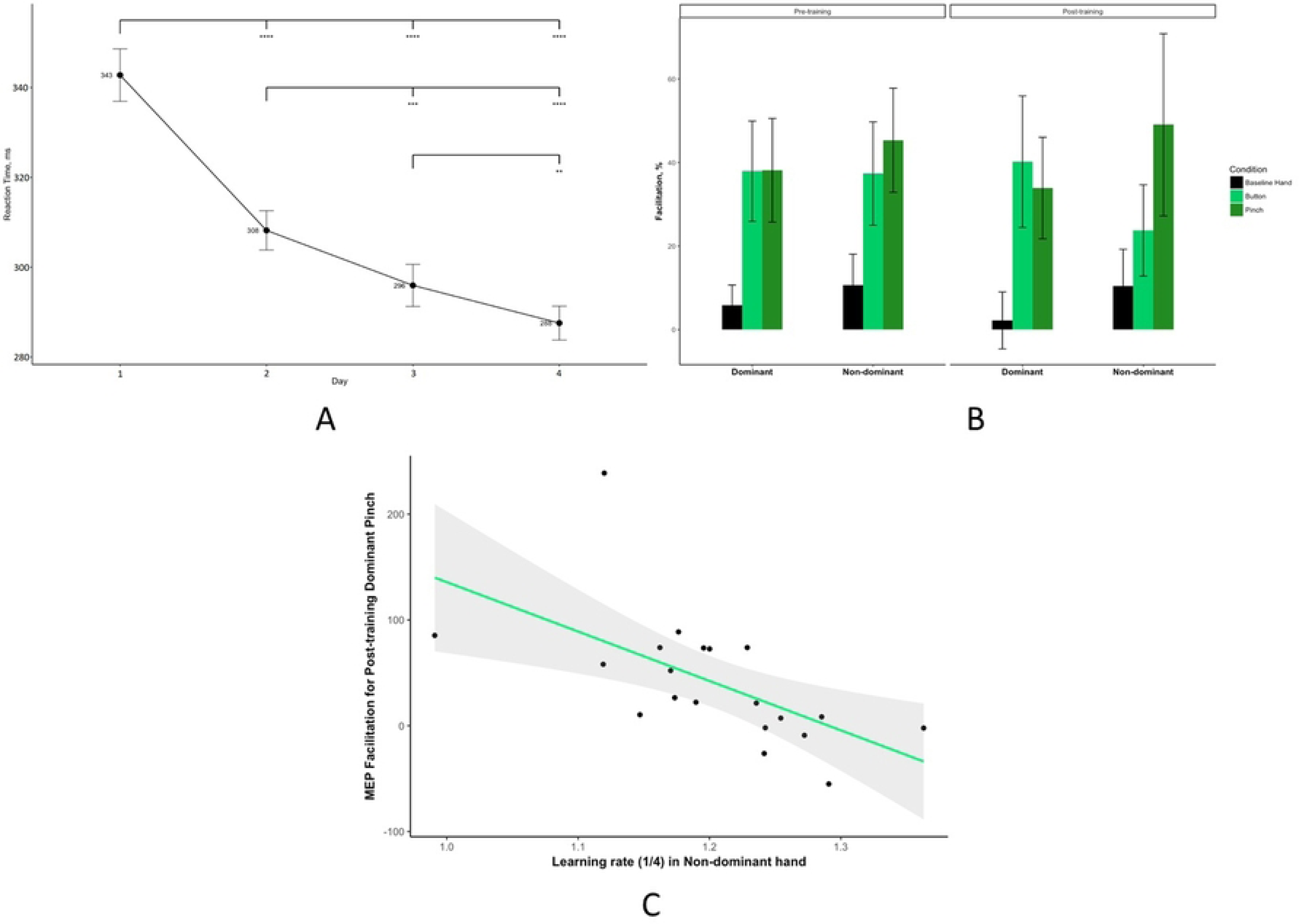
A) Results of the SRT task. Mean RT (in ms) is presented for each of the four days of the training. Black circles depict the mean RT for each day. Error bars depict the standard error of the mean (** – p < .01, *** – p < .001, **** – p < .0001). B) Changes in MEP amplitude compared with baseline (in %) during the observation of the static hand (Baseline Hand – black), observation of the button-pressing movement (Button – light green), and observation of the pinch-to-grip movement (Pinch – dark green) in the dominant and nondominant hemispheres (on the x-axis) on the pretraining (left) and post-training (right) sessions. Data presented as mean ± standard error of mean. C) Scatter plot depicting the association between the learning rate in the nondominant hand (on the x-axis) and MEP facilitation during the observation of pinch-to-grip movement in the dominant hand in the post-training session (on the y-axis). The green line represents the fitted linear function of the data, and the gray area represents the 95% confidence level interval.

### MEP estimation

#### Comparison of baselines

The main dependent variable for this phase of the experiment was MEP peak-to-peak amplitude in the FDI muscle. However, assessing MEP amplitudes in ADM was also important to ensure that we observed muscle-specific changes in corticospinal excitability, truly reflecting the mirror effect. First, three-way repeated measures ANOVA was performed only for the pre- and postobservation baseline data (2 (Condition: Baseline or Baseline Post) x 2 (Hand: Dominant or Nondominant) x 2 (Session: Pretraining or Post-training)) for raw MEP data for both muscles to ensure the absence of general corticospinal facilitation effect inside the TMS sessions. Because all the independent variables had only two levels, Mauchly’s test was not performed. This analysis showed neither significant main effects nor significant interaction effects for both muscles. Most importantly, the main effect of Condition was significant neither for FDI (F(1, 21) = 0.02, p = .878, h^2^ = .00001), nor for ADM (F(1, 21) = 1.65, p = .212, h^2^ = .001). Therefore, for the following analysis, the Baseline and Baseline Postconditions were collapsed, forming a single baseline measurement.

Because we primarily aimed to investigate motor resonance, which is represented by an increase in corticospinal excitability during action observation, for the subsequent analysis, we have calculated the ratio of MEP amplitudes in the observation conditions to the collapsed baseline condition for each muscle. As a result, we have obtained the new dependent variable—change (facilitation) of the corticospinal excitability (in percentage)—in the three remaining conditions: Baseline Hand, Button, and Pinch. This calculation was performed for both muscles.

#### Difference in mirror effect between sessions

Assessment of changes in the facilitation of corticospinal excitability in FDI muscle was carried out with the help of a three-way repeated measures ANOVA: 3 (Condition: Baseline Hand, Button, or Pinch) x 2 (Session: Pretraining or Post-training) x 2 (Hemisphere: Dominant or Nondominant). Mauchly’s test showed no violations of the sphericity assumption, so corrections were not implemented. This analysis revealed only the significant main effect of Condition (F(2.44) = 8.98, p < .001, h^2^ = .062), while all the other effects, and, most importantly, the main effect of Session (F(1, 22) = 0.284, p = .599, h^2^ = .0005) and the interaction effect between Session and Condition (F(2, 44) = 0.11, p = .89, h^2^ = .0004), were not significant. As a result, one-way repeated measures ANOVA was performed on the data collapsed for sessions and hemispheres to assess the main effect of condition. This analysis, as expected, demonstrated a significant main effect (F(2, 44) = 8.98, p < .001, h^2^ = .112). Visualization of MEP facilitation across different conditions, sessions, and hemispheres is presented in Fig 2b.

#### Mirror effect

To investigate the revealed main effect of the condition in greater detail and check whether the mirror effect in general is present, post hoc comparisons were made. For this, a paired-samples t-test with Holm–Bonferroni correction was used. Facilitation of the corticospinal excitability during observation of the static hand (M = 7.23%, SD = 33.78%) was significantly different from the facilitation during observation of the button-pressing movement (M = 34.80%, SD = 61.10%) (t(22) = 3.38, p < .01) and observation of the pinch-to-grip movement (M = 41.60%, SD = 72.23%) (t(22) = 3.31, p < .01). The difference in the facilitation of excitability during observation of the button-pressing movement and pinch-to-grip movement was not significant (t(22) = 1.00, p = .33). Facilitation data for conditions only (merged across sessions and hemispheres) are depicted in Supplementary Fig 1.

#### Changes in MEP amplitudes in the ADM muscle

In the following step of the analysis, changes in the corticospinal excitability of the ADM muscle were assessed. For this step, a three-way repeated measures ANOVA was used: 3 (Condition: Baseline Hand, Button, or Pinch) x 2 (Session: Pretraining or Post-training) x 2 (Hand: Dominant or Nondominant)). This analysis showed the absence of any significant effects, main or interaction alike. Most importantly, the main effect of the Condition was not significant (F(2, 44) = 1.29, p = .285, h^2^ = .0066), demonstrating that the facilitation of MEP amplitude does not change across conditions for the ADM muscle. Based on this, any effect we observe only in the FDI muscle (target muscle) most likely reflects the activity of MNS because no changes in the ADM muscle (control muscle) were found, supporting the specificity of potential effects.

### Analysis of correlation

Study of the relationship between the speed of sensorimotor learning and the levels of motor resonance in pre-training and post-training sessions (Fig 2c). The ratio of mean RT on the first day of the behavioral task to mean RT on the last day of the behavioral task was used as an index of the sensorimotor learning rate. This was done separately for the dominant and nondominant hands. Overall, 16 correlations were calculated: values of motor facilitation in both conditions (during observation of pinch-to-grip and button-pressing movements) in two hemispheres during two sessions (12 variables) were compared with the learning rates in two hands (two variables). The Bonferroni method adjusted the p-values and accounted for the multiple comparisons issue. After this adjustment was implemented, no correlations were significant.

## Discussion

The present study explored sensorimotor learning and its relationship with motor resonance and mirror neuron activity. The SRT task was employed over four days, leading to a significant decrease in RT, hence indicating successful sensorimotor learning. Observation of button-pressing and pinch-to-grip movements induced mirror neuron activity, as evidenced by MEP facilitation. However, no significant correlation was found between sensorimotor learning and changes in corticospinal excitability during action observation, challenging the proposed connection between motor resonance and learning.

### Sensorimotor Learning

Analysis of SRT task results revealed a significant decrease in the mean RT across the four days of training, which indicates that the implemented design was sufficient for inducing sensorimotor learning in the participants. Even though the main effect of Hand on RT was also significant, the absence of a significant interaction effect signifies that the learning rate itself was not different between the two hands. Altogether, these data suggest that learning did occur as a result of the completion of the implemented SRT task, putatively inducing a sensorimotor association between button-pressing movement and simple geometric stimuli (green circles on the screen), and that rates of this type of learning were not different between dominant and nondominant hands.

It is possible that performance on the SRT task was generally better for the dominant hand because of control asymmetries that can affect adaptive processes. It has previously been shown that there is no difference between the speed or final degree of adaptation for both hands, suggesting that visuomotor adaptation possibly occurs similarly for the two hands (16).

### Mirror effect

Post hoc comparisons, implemented on the electromyography data from FDI muscle collapsed across Hands and Sessions, demonstrated that MEP facilitation was significantly different for the observation of button-pressing and pinch-to-grip movements compared with the observation of the static hand. This effect suggests that the implemented setup was adequate for eliciting motor resonance (mirror effect), manifesting in a muscle-specific increase of corticospinal excitability during observation of the movement performed by another individual. The muscle specificity of this effect is ensured by the absence of any significant differences across conditions for MEP facilitation in the ADM muscle.

Mirror neurons are activated when an action is observed in areas of the cerebral cortex responsible for the muscles involved in the viewed activity (17–19). Thus, our results are in agreement with a large body of literature on this topic.

### Correlation between sensorimotor learning and MEP facilitation

Our first hypothesis stated that sensorimotor learning in the SRT task would be correlated with the facilitation of corticospinal excitability during action observation in the pretraining session, reflecting a connection of the motor resonance phenomenon with the sensorimotor learning processes. However, all estimated correlations were not significant not only for the pretraining session, but also for the post-training session, thus demonstrating the absence of an association between sensorimotor learning rate and changes in corticospinal excitability during action observation in this particular experimental setup. This is inconsistent with the literature. Studies have shown that changes in the excitability of the corticospinal system are the result of learning to move (20–22). Perhaps the differences in the results are because, when studying changes in the activity of the corticospinal system after learning to move, TMS is used during the movement, not during the observation of the movement. Moreover, our second hypothesis, according to which we expected to observe changes in the pattern of MEP facilitation during action observation induced by the process of sensorimotor learning in the SRT task, was also not supported by the results of our experiment. Analysis of variance carried out on the MEP facilitation data for the FDI muscle revealed the absence of any significant effects apart from the main effect of the condition. Therefore, the pattern of action observation–induced changes in corticospinal excitability did not differ between the pretraining and post-training sessions. In other words, our results do not support the ASL theory, according to which the motor resonance phenomenon, as a manifestation of mirror neurons’ activity, is developed through sensorimotor learning and can be modulated through this process. These results were unexpected because the studies in this field have demonstrated that the motor resonance phenomenon could be modulated in the process of sensorimotor learning (6,23–25). However, certain differences in the experimental design might have contributed to the observed discrepancies between our results and the data in the field. In previous studies, sensorimotor learning was based on the nonmatching action observation stimuli (learning to perform index-finger abduction in response to the observation of little-finger abduction), while in our experiment, training was based on the novel simple geometric stimuli (green circles on the screen), thus bearing a purely additive nature and not causing a mismatch in existing sensorimotor matching between observed and performed actions. One recent relevant study, in which the coupling of performed actions with observed simple geometric stimuli was induced, was carried out with the help of fMRI without assessment of motor response itself, thus making it problematic to compare it with the outcomes of our study (26). Moreover, in our study, long-term training (four days) was implemented, comparing two training sessions in existing studies (6,23,24). However, more research is needed to establish whether this contradiction arises because of the discrepancies in experimental design or the nature of the studied phenomena per se.

Although we have not confirmed ASL theory, we have demonstrated that the rate of nondominant hand learning correlates with the activation of mirror neurons in the dominant hemisphere after training in button pressing and pinching. In addition, the learning rate of the dominant hand correlates with the response of mirror neurons in the nondominant hemisphere when observing the baseline hand.

Our findings may be related to the lateralization of motor learning. The model proposed by Pratik K. Mutha and colleagues suggests that the left hemisphere provides predictive control mechanisms that determine certain aspects, such as the direction of movement for movements of both the contralateral and ipsilateral arms. At the same time, the right hemisphere also contributes to positional control mechanisms during the movements of either arm (16).

This separation of functions may be a way to ensure that none of the processes are completely compromised—the dominant hemisphere can ensure the precise execution of a task. In contrast, the nondominant hemisphere gradually learns to improve movement planning.

Thus, the correlation between nondominant hand learning and dominant hemisphere mirroring neuron activation may result from nondominant hemisphere learning to improve movement planning (16).

Also, the presence of this ipsilateral interaction observed through the correlation between nondominant hand learning and mirror neuron activity in the dominant hemisphere after learning may be related to the ipsilateral component of the corticospinal system. The function of this is currently not known.

Neuroimaging studies have revealed activity in the precentral gyrus ipsilateral to the side of hand movement, especially during complex finger movements, in healthy adults (27) and after stroke recovery (27,28). In addition, TMS can cause hand movements ipsilateral to the stimulated hemisphere (29). The precentral gyrus areas of the ipsilateral side of the arm movement were interpreted as part of the primary motor areas or Brodmann area 4. This interpretation was based on studies in nonhuman primates that clearly demonstrated ipsilateral or bilateral motor representations within the primary motor areas (30).

The ipsilateral interaction shown in the present study is consistent with the results of previous fMRI studies in the ventral premotor areas involved in distal finger movements. These studies show that accurate grip performance is associated with stronger signaling in the inferior frontal gyrus, ipsilateral to the responsive hand (31), and that the ventral premotor cortex has ipsilateral representations of the fingers (27,32).

Thus, although the present study refutes ASL theory, it also shows that there is a correlation between nondominant hand learning and mirror neuron activation in the dominant hemisphere. In the literature, it has been demonstrated that ipsilateral interaction for the hands exists and is most likely associated with the corticospinal system.

## Conclusion

In the current study, we have investigated whether the degree of motor resonance during action observation can act as a predictor of the sensorimotor learning rate and how sensorimotor learning modulates the facilitation of corticospinal excitability during action observation. We observed a statistically significant decrease of RTs in the process of learning in the SRT task and differences in motor facilitation during action observation compared with the observation of the static hand, thus reflecting the process of putative mirror neurons’ activity. However, our data have demonstrated that the sensorimotor learning rate is not associated with either pretraining or post-training estimates of motor facilitation during action observation and that sensorimotor learning does not affect the pattern of motor resonance. Therefore, the outcome of our study does not support ASL theory and does not show any significant association between the process of sensorimotor learning and mirror neurons’ activity being reflected in the motor resonance phenomenon.

Our findings have broader implications for understanding the complexities of motor learning and intricate interplay between the brain’s mirror neuron system and sensorimotor processes. Although our results did not support ASL theory’s proposed connection between sensorimotor learning and mirror neuron activity, they challenge existing assumptions, calling for a reevaluation of the underlying mechanisms driving motor resonance. The present study contributes to a growing body of literature seeking to unravel the multifaceted nature of sensorimotor learning and mirror neuron functioning. By examining the correlation between sensorimotor learning rates and the motor resonance phenomenon, we have provided valuable insights into the dynamic interplay between these processes. This knowledge can inform future research and guide the development of innovative approaches to motor rehabilitation and cognitive interventions.

The limitations of our study, such as the specific experimental design and focus on a novel set of stimuli, highlight the need for further investigations to elucidate the complex relationship between sensorimotor learning, mirror neuron activity, and the modulation of motor resonance. Future studies employing diverse methodologies, larger sample sizes, and different learning paradigms are warranted to expand our understanding of these interconnected phenomena.

## Acknowledgments

This work is an output of a research project implemented as part of the Basic Research Program at the National Research University Higher School of Economics (HSE University) and was carried out using HSE Automated system of non-invasive brain stimulation with the possibility of synchronous registration of brain activity and registration of eye movements

## Supporting information

**Supplementary Fig 1.**
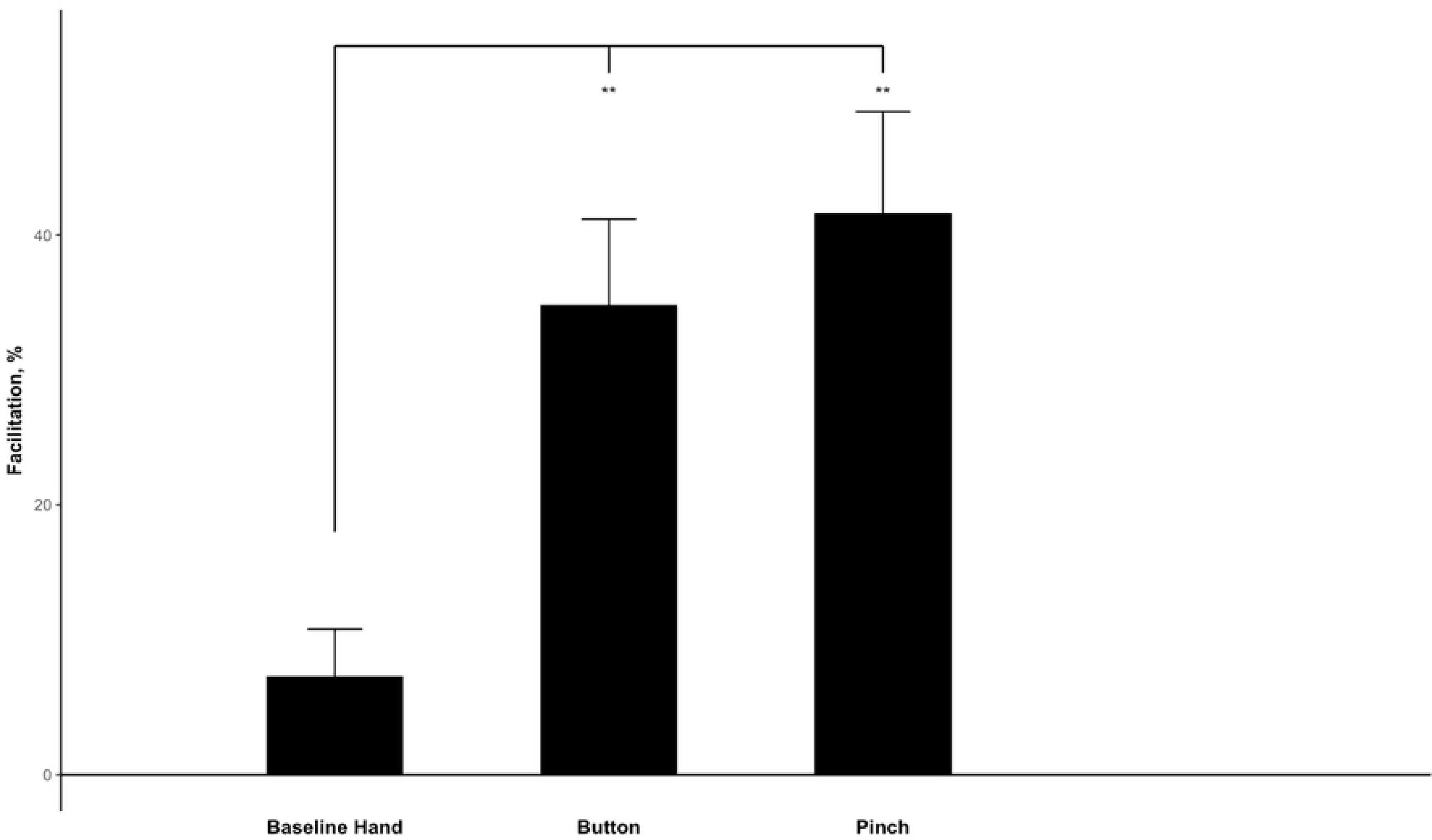
Changes in MEP amplitude in the nondominant hemispheres compared with baseline (in %) during the observation of the static hand (Baseline Hand), observation of the button-pressing movement (Button), and observation of the pinch-to-grip movement (Pinch). Data are presented as mean ± standard error of mean (** – p < .01)

## References

1. di Pellegrino G, Fadiga L, Fogassi L, Gallese V, Rizzolatti G. Understanding motor events: a neurophysiological study. Exp Brain Res [Internet]. 1992 Oct [cited 2023 Aug 23];91(1):176–80. Available from: https://pubmed.ncbi.nlm.nih.gov/1301372/

2. Kilner JM, Marchant JL, Frith CD. Relationship between Activity in Human Primary Motor Cortex during Action Observation and the Mirror Neuron System. PLoS One [Internet]. 2009 Mar 17 [cited 2023 Aug 23];4(3):e4925. Available from: https://journals.plos.org/plosone/article?id=10.1371/journal.pone.0004925

3. Mukamel R, Ekstrom AD, Kaplan J, Iacoboni M, Fried I. Single-neuron responses in humans during execution and observation of actions. Curr Biol [Internet]. 2010 Apr 27 [cited 2023 Aug 23];20(8):750–6. Available from: https://pubmed.ncbi.nlm.nih.gov/20381353/

4. Meltzoff AN, Moore MK. Imitation in Newborn Infants: Exploring the Range of Gestures Imitated and the Underlying Mechanisms. Dev Psychol [Internet]. 1989 [cited 2023 Aug 23];25(6):954–62. Available from: https://pubmed.ncbi.nlm.nih.gov/25147405/

5. Catmur C. Sensorimotor learning and the ontogeny of the mirror neuron system. Neurosci Lett [Internet]. 2013 Apr 12 [cited 2023 Aug 23];540:21–7. Available from: https://pubmed.ncbi.nlm.nih.gov/23063950/

6. Catmur C, Walsh V, Heyes C. Sensorimotor learning conFigs the human mirror system. Curr Biol [Internet]. 2007 Sep 4 [cited 2023 Aug 23];17(17):1527–31. Available from: https://pubmed.ncbi.nlm.nih.gov/17716898/

7. Heyes C. Where do mirror neurons come from? Neurosci Biobehav Rev [Internet]. 2010 Mar [cited 2023 Aug 23];34(4):575–83. Available from: https://pubmed.ncbi.nlm.nih.gov/19914284/

8. Naish KR, Houston-Price C, Bremner AJ, Holmes NP. Effects of action observation on corticospinal excitability: Muscle specificity, direction, and timing of the mirror response. Neuropsychologia [Internet]. 2014 Nov 1 [cited 2023 Aug 23];64:331–48. Available from: https://pubmed.ncbi.nlm.nih.gov/25281883/

9. Catmur C, Walsh V, Heyes C. Associative sequence learning: the role of experience in the development of imitation and the mirror system. Philos Trans R Soc Lond B Biol Sci [Internet]. 2009 Aug 27 [cited 2023 Aug 23];364(1528):2369–80. Available from: https://pubmed.ncbi.nlm.nih.gov/19620108/

10. Keysers C, Gazzola V. Hebbian learning and predictive mirror neurons for actions, sensations and emotions. Philos Trans R Soc Lond B Biol Sci [Internet]. 2014 Jun 5 [cited 2023 Aug 23];369(1644). Available from: https://pubmed.ncbi.nlm.nih.gov/24778372/

11. Keysers C, Perrett DI. Demystifying social cognition: a Hebbian perspective. Trends Cogn Sci [Internet]. 2004 [cited 2023 Aug 23];8(11):501–7. Available from: https://pubmed.ncbi.nlm.nih.gov/15491904/

12. Shin YK, Proctor RW, Capaldi EJ. A review of contemporary ideomotor theory. Psychol Bull [Internet]. 2010 Nov [cited 2023 Aug 23];136(6):943–74. Available from: https://pubmed.ncbi.nlm.nih.gov/20822210/

13. Cracco E, Bardi L, Desmet C, Genschow O, Rigoni D, Coster L De, et al. Automatic imitation: A meta-analysis. Psychol Bull [Internet]. 2018 May 1 [cited 2023 Aug 23];144(5):453–500. Available from: https://pubmed.ncbi.nlm.nih.gov/29517262/

14. Rossi S, Hallett M, Rossini PM, Pascual-Leone A, Avanzini G, Bestmann S, et al. Safety, ethical considerations, and application guidelines for the use of transcranial magnetic stimulation in clinical practice and research. Clin Neurophysiol [Internet]. 2009 Dec [cited 2023 Aug 23];120(12):2008–39. Available from: https://pubmed.ncbi.nlm.nih.gov/19833552/

15. Rossini PM, Burke D, Chen R, Cohen LG, Daskalakis Z, Di Iorio R, et al. Non-invasive electrical and magnetic stimulation of the brain, spinal cord, roots and peripheral nerves: Basic principles and procedures for routine clinical and research application. An updated report from an I.F.C.N. Committee. Clin Neurophysiol [Internet]. 2015 Jun 1 [cited 2023 Aug 23];126(6):1071–107. Available from: https://pubmed.ncbi.nlm.nih.gov/25797650/

16. Mutha PK, Haaland KY, Sainburg RL. THE EFFECTS OF BRAIN LATERALIZATION ON MOTOR CONTROL AND ADAPTATION. J Mot Behav [Internet]. 2012 [cited 2023 Aug 22];44(6):455. Available from: /pmc/articles/PMC3549328/

17. Wurm MF, Lingnau A. Decoding Actions at Different Levels of Abstraction. The Journal of Neuroscience [Internet]. 2015 May 5 [cited 2023 Aug 23];35(20):7727. Available from: /pmc/articles/PMC6795187/

18. Wurm MF, Caramazza A. Distinct roles of temporal and frontoparietal cortex in representing actions across vision and language. Nat Commun [Internet]. 2019 Dec 1 [cited 2023 Aug 23];10(1). Available from: /pmc/articles/PMC6336825/

19. Heyes C, Catmur C. What Happened to Mirror Neurons? Perspectives on Psychological Science [Internet]. 2022 Jan 1 [cited 2023 Aug 23];17(1):153. Available from: /pmc/articles/PMC8785302/

20. Koeneke S, Lutz K, Herwig U, Ziemann U, Jäncke L. Extensive training of elementary finger tapping movements changes the pattern of motor cortex excitability. Exp Brain Res [Internet]. 2006 Sep [cited 2023 Aug 23];174(2):199–209. Available from: https://pubmed.ncbi.nlm.nih.gov/16604315/

21. Bagce HF, Saleh S, Adamovich S V., Krakauer JW, Tunik E. Corticospinal excitability is enhanced after visuomotor adaptation and depends on learning rather than performance or error. J Neurophysiol [Internet]. 2013 Feb 1 [cited 2023 Aug 23];109(4):1097. Available from: /pmc/articles/PMC3569119/

22. Cirillo J, Todd G, Semmler JG. Corticomotor excitability and plasticity following complex visuomotor training in young and old adults. Eur J Neurosci [Internet]. 2011 Dec [cited 2023 Aug 23];34(11):1847–56. Available from: https://pubmed.ncbi.nlm.nih.gov/22004476/

23. Catmur C, Mars RB, Rushworth MF, Heyes C. Making mirrors: premotor cortex stimulation enhances mirror and counter-mirror motor facilitation. J Cogn Neurosci [Internet]. 2011 Sep [cited 2023 Aug 23];23(9):2352–62. Available from: https://pubmed.ncbi.nlm.nih.gov/20946056/

24. Cavallo A, Heyes C, Becchio C, Bird G, Catmur C. Timecourse of mirror and counter-mirror effects measured with transcranial magnetic stimulation. Soc Cogn Affect Neurosci [Internet]. 2014 Aug 1 [cited 2023 Aug 23];9(8):1082–8. Available from: https://pubmed.ncbi.nlm.nih.gov/23709352/

25. Taschereau-Dumouchel V, Hétu S, Michon PE, Vachon-Presseau E, Massicotte E, De Beaumont L, et al. BDNF Val66Met Polymorphism Influences Visuomotor Associative Learning and the Sensitivity to Action Observation. Sci Rep [Internet]. 2016 Oct 5 [cited 2023 Aug 23];6. Available from: https://pubmed.ncbi.nlm.nih.gov/27703276/

26. Press C, Catmur C, Cook R, Widmann H, Heyes C, Bird G. FMRI evidence of “mirror” responses to geometric shapes. PLoS One [Internet]. 2012 Dec 14 [cited 2023 Aug 23];7(12). Available from: https://pubmed.ncbi.nlm.nih.gov/23251653/

27. Cramer SC, Finklestein SP, Schaechter JD, Bush G, Rosen BR. Activation of distinct motor cortex regions during ipsilateral and contralateral finger movements. J Neurophysiol [Internet]. 1999 [cited 2023 Aug 22];81(1):383–7. Available from: https://pubmed.ncbi.nlm.nih.gov/9914297/

28. Weiller C, Chollet F, Friston KJ, Wise RJS, Frackowiak RSJ. Functional reorganization of the brain in recovery from striatocapsular infarction in man. Ann Neurol [Internet]. 1992 [cited 2023 Aug 23];31(5):463–72. Available from: https://pubmed.ncbi.nlm.nih.gov/1596081/

29. Wassermann EM, Pascual-Leone A, Hallett M. Cortical motor representation of the ipsilateral hand and arm. Exp Brain Res [Internet]. 1994 Jul [cited 2023 Aug 23];100(1):121–32. Available from: https://link.springer.com/article/10.1007/BF00227284

30. Gentilucci M, Fogassi L, Luppino G, Matelli M, Camarda R, Rizzolatti G. Somatotopic representation in inferior area 6 of the macaque monkey. Brain Behav Evol [Internet]. 1989 [cited 2023 Aug 23];33(2–3):118–21. Available from: https://pubmed.ncbi.nlm.nih.gov/2758288/

31. Ehrsson HH, Fagergren A, Jonsson T, Westling G, Johansson RS, Forssberg H. Cortical activity in precision-versus power-grip tasks: an fMRI study. J Neurophysiol [Internet]. 2000 [cited 2023 Aug 22];83(1):528–36. Available from: https://pubmed.ncbi.nlm.nih.gov/10634893/

32. Hanakawa T, Parikh S, Bruno MK, Hallett M. Finger and face representations in the ipsilateral precentral motor areas in humans. J Neurophysiol [Internet]. 2005 May [cited 2023 Aug 22];93(5):2950. Available from: /pmc/articles/PMC1440886/

